# Investigating the unique ability of *Trichodesmium* to fix carbon and nitrogen simultaneously using MiMoSA

**DOI:** 10.1101/2022.10.14.512281

**Authors:** Joseph J. Gardner, Bri-Mathias S. Hodge, Nanette R. Boyle

## Abstract

The open ocean is an extremely competitive environment, partially due to the dearth of nutrients. *Trichodesmium erythraeum*, a marine diazotrophic cyanobacterium, is a keystone species in the ocean due to its ability to fix nitrogen and leak 30-50% into the surrounding environment, providing a valuable source of a necessary macronutrient to other species. While there are other diazotrophic cyanobacteria that play an important role in the marine nitrogen cycle, *Trichodesmium* is unique in its ability to fix both carbon and nitrogen simultaneously during the day without the use of specialized cells called heterocysts to protect nitrogenase from oxygen. Here, we use the advanced modeling framework called <u>M</u>ultiscale <u>M</u>ulti<u>o</u>bjective <u>S</u>ystems <u>A</u>nalysis (MiMoSA) to investigate how *Trichodesmium erythraeum* can reduce dimolecular nitrogen to ammonium in the presence of oxygen. Our simulations indicate that nitrogenase inhibition is best modeled as Michealis Menten competitive inhibition and that cells along the filament maintain microaerobia using high flux through Mehler’s reactions in order to protect nitrogenase from oxygen. We also examined the effect of location on metabolic flux and found that cells at the end of filaments operate in distinctly different metabolic modes than internal cells despite both operating in a photoautotrophic mode. These results give us important insight into how this species is able to operate photosynthesis and nitrogen fixation simultaneously, giving it a distinct advantage over other diazotrophic cyanobacteria because they can harvest light directly to fuel the energy demand of nitrogen fixation.

**IMPORTANCE:** *Trichodesmium erythraeum* is a marine cyanobacterium responsible for approximately half of all biologically fixed nitrogen, making it an integral part of the global nitrogen cycle. Interestingly, unlike other nitrogen fixing cyanobacteria, *Trichodesmium* does not use temporal or spatial separation to protect nitrogenase from oxygen poisoning; instead, it operates photosynthesis and nitrogen fixation reactions simultaneously during the day. Unfortunately, the exact mechanism the cells utilize to operate carbon and nitrogen fixation simultaneously is unknown. Here, we use an advanced metabolic modeling framework to investigate and identify the most likely mechanisms *Trichodesmium* uses to protect nitrogenase from oxygen. The model predicts that cells operate in a microaerobic mode, using both respiratory and Mehler reactions to dramatically reduce intracellular oxygen concentrations.

## INTRODUCTION

*Trichodesmium erythraeum*, a filamentous diazotrophic cyanobacterium, is a keystone organism in marine environments due to its significant contribution to the global nitrogen cycle. The *Trichodesmium* genus is responsible for approximately 49% of the reactive nitrogen produced in the ocean every year (4- 8). What makes this organism even more intriguing is its ability to simultaneously fix carbon and nitrogen, making it an important contributor in two global macronutrient cycles. The ability to fix both carbon and nitrogen during daylight hours stems from a unique approach to regulating these processes within the filament. While much is still unknown about the exact mechanism of how *Trichodesmium* regulates intracellular oxygen during the day without structural elements to protect nitrogenase, what is known is discussed below. Along the filament, cells are divided into two distinct groups: nitrogen-fixing (diazotrophic) cells and carbon-fixing (photoautotrophic) cells (7). Diazotrophic cells are grouped together in the center of the filament, often in one continuous section but can be found in up to three sections of varying cell length, forming diazocyte regions. The balance of the filament is made up of photoautotrophic cells, which have been reported to be between 70 – 85% of the population (9). The two different cell types exchange metabolites back and forth in a symbiotic relationship. Diazotrophs fix dimolecular nitrogen into biologically reactive compounds such as ammonium, urea, and L-aspartate-L-arginine chains called cyanophycin, and provide these nitrogen rich compounds to photoautotrophs in exchange for reduced carbon (glucose, glycogen, etc.). Individual cells are therefore responsible not only for their growth, but for the growth of other cells within the trichome and the whole community which creates an effective division of labor between cell types. The presence of two distinct cell populations makes accurate modeling of the growth of *Trichodesmium* difficult. Yet another challenge is the dynamic nature of the growth of *Trichodesmium*. Unlike some cyanobacterial species, which can grow in continuous light without adverse effect on cellular physiology, *T. erythraeum* requires diel light cycles in order to thrive, and with these diel cycles comes large changes in biomass composition and cellular objectives (2). When grown in constant light, nitrogenase enzyme activity declines and cells are unable to support growth (10) without nitrogen-supplemented medium. While the specific roles of the day versus night period have not been formally described, evidence suggests that communities maximize metabolite acquisition during the day – similar to other unicellular photosynthetic marine organisms – and prepare the colony for metabolite acquisition at night (10-12). Additionally, metabolite acquisition requires a fine balance in diazotrophic photosynthetic organisms: photosystem II (PSII) releases O_2_, which irreversibly inhibits nitrogenase. *T. erythraeum* partially compensates for potential inhibition by downregulating PSII in diazotrophic cells, but oxygen can readily diffuse through cellular membranes without the same structural elements heterocystic cyanobacteria use to prevent oxygen diffusion to nitrogen fixing cells (13). In contrast to temporal regulation – through nitrogenase activity in the dark – or specialized O_2_- excluding compartment formation (14), we hypothesize that *T. erythraeum* primarily uses metabolic means to protect nitrogenase from O_2_. *T. erythraeum* can consume O_2_ through Mehler reactions, which reverses O_2_ production by PSII through hydrogen peroxide biosynthesis, enabling the cell to protect nitrogenase from oxygen with minimal energy input (15-18). Evidence suggests that glycolysis energetically drives diazotrophy instead of light (16, 17), creating a strong link between carbon and nitrogen metabolism (19, 20). This is augmented by robust L-arginine degradation pathways that allow for organic nitrogen compounds to be used as nitrogen, energy, and carbon sources. Given the complexity of the growth and metabolism of *Trichodesmium*, simple metabolic models such as flux balance analysis(21) and its many augmented forms(22-24) cannot adequately describe the organism. Instead, we have developed a framework, called Multiscale Multiobjective Systems Analysis (MiMoSA)(2)(see Figure 1), that is capable of tracking individual cells and nutrients in space and time and predict cell differentiation and metabolic fluxes. The MiMoSA framework is also able to model the heterogeneity that occurs in the system due to the different cell types and spatial distribution of nitrogen compounds. The MiMoSA framework takes in data on the cell in the form of the genome-scale metabolic model and associated constraints for each cell type, the environmental conditions (light, CO_2_ percent, N_2_ percent, presence of exogenous nitrogen, etc.), the initial cell conditions (filament lengths, relative quantities of cells), and the rules by which cells will reduce O_2_ and regulate nitrogen fixation in the presence of other sources. The model is then able to simulate growth rates, individual intracellular fluxes, and cell and nutrient tracking. This modeling framework is specifically able to determine how spatiotemporal organizations of two cell types interact with the environment to manifest the phenotypes observed in culture. Here, we present an updated version of the MiMoSA model for *Trichodesmium* which enumerates all the mechanisms for degradation and assimilation of L-arginine. The updated model was used to further investigate the exact mechanisms of how *Trichodesmium* is able to carry out simultaneous carbon and nitrogen fixation.

**Figure 1.**
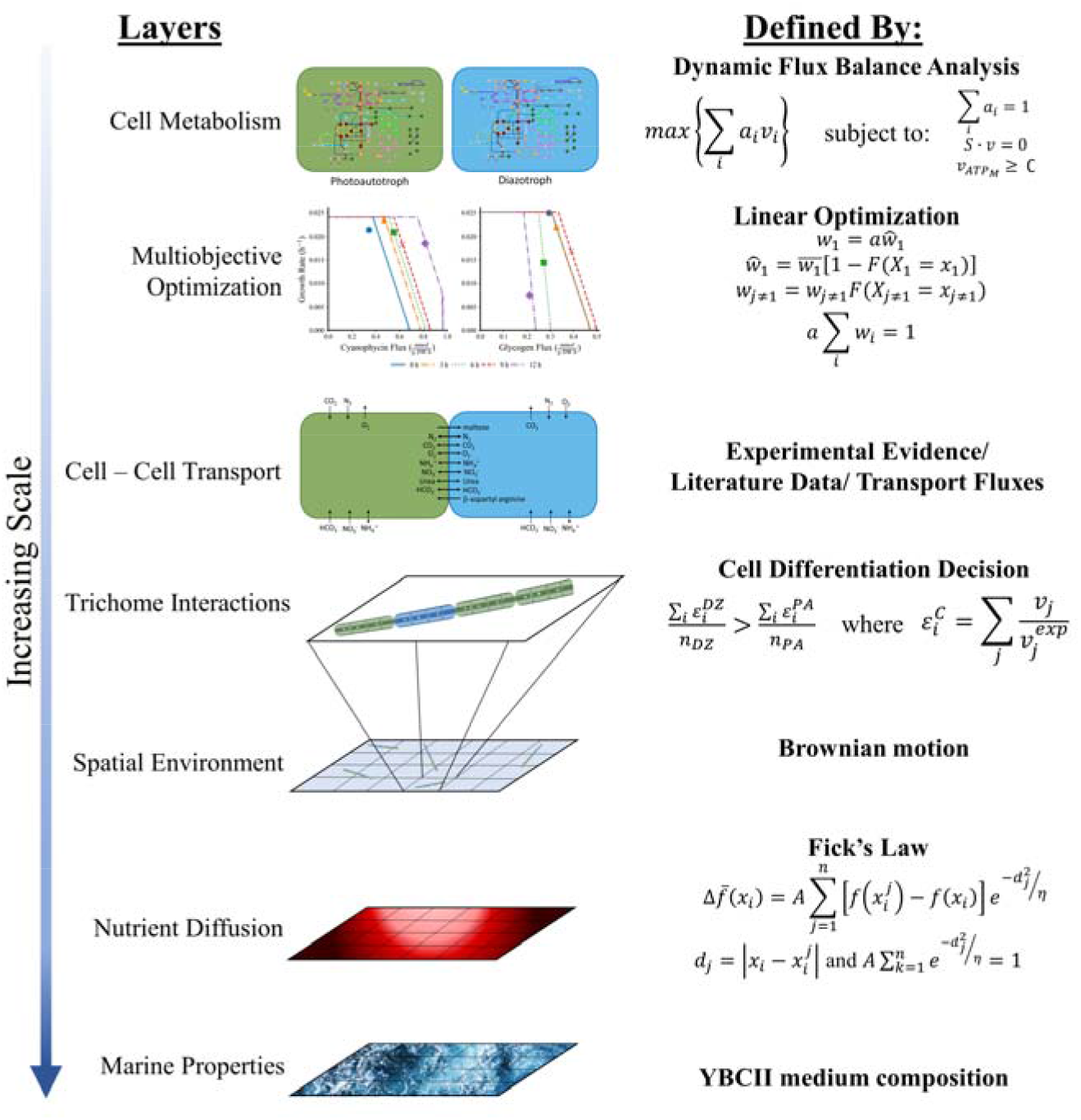
Overview of the MiMoSA framework. The MiMoSA modeling framework is made up of three agents: cells, filaments and ocean. The cells agent has two subclasses: photoautotroph and diazotroph. Each cell is represented by an individual agent in the model that can act independently. Intracellular metabolic fluxes are modeled using dynamic flux balance analysis with multiobjective optimization. There is a filament agent, which defines the intracellular transport and determines whether new cells are differentiated into a photoautotoph or diazotroph based on the need for carbon or nitrogen in the filament. Cells are randomly seeded in the simulation and are allowed to move via Brownian motion. Finally, the last agent in the framework is the ocean agent, which tracks nutrient concentrations in space and time. For a detailed description of the equations describing each of these processes, please refer to the original MiMoSA model(2).

## MATERIALS & METHODS

### Updated Genome-Scale Metabolic Model

We updated the published genome-scale metabolic network reconstruction of *T. erythraeum* (25) to include an additional 327 reactions and 157 metabolites and replace18 diffusion-based transport reactions with more informed transport reactions (such as ABC, passive, coupled, or other transporters) increasing its size to include a total of 1,293 reactions and 1,147 metabolites. The updated reactions, associated genes, and metabolites are provided as Supplemental File 1, the complete genome-scale metabolic network is provided as Supplemental File 2, and the model with bounds updated to allow biomass catabolism is provided as Supplemental File 3.

### Multiscale Multiobjective Systems Analysis (MiMoSA) Framework

The previously described MiMoSA framework (shown in Figure 1) (2) was updated to reformulate the multiobjective optimization process using methods described by Zomorrodi et al. (26, 27) to solve a bilevel objective for each individual cell (as opposed to an individual cell subject to the community optimization). For the model of *Trichodesmium*, the *a priori* objective (the combinatorial synthesis of biomass and the transactional metabolite, either β-aspartyl arginine or maltose) is optimized first and then becomes a constraint for the second optimization of metabolite storage. The community fitness is still predicated on the summary behavior of completely individually optimized cells. Cells, however, do have a choice between anabolism, primary production, and catabolism, prioritizing them respectively. We also updated how metabolites are allowed to diffuse by including additional compartments to differentiate the free diffusion within the filament (“_f”) from non-diffusing biomass compounds (“_b”), extracellular compounds (“_e”), and cytosolic compounds (“_c”) as shown in Figure 2. The updated and complete MiMoSA framework is provided as two files: the Python modules are in Supplemental File 4 and the Agent-Based model in Supplemental File 5. Algorithms for different growth regimes (anabolism, catabolism, and primary productivity), as well as the steps cells undertake in optimizing metabolism, are in Supplemental Files 6 – 9. A complete list of allowable transport between compartments, cell, and the environment are provided in Supplemental File 10. *Objective Equation Selection*

**Figure 2.**
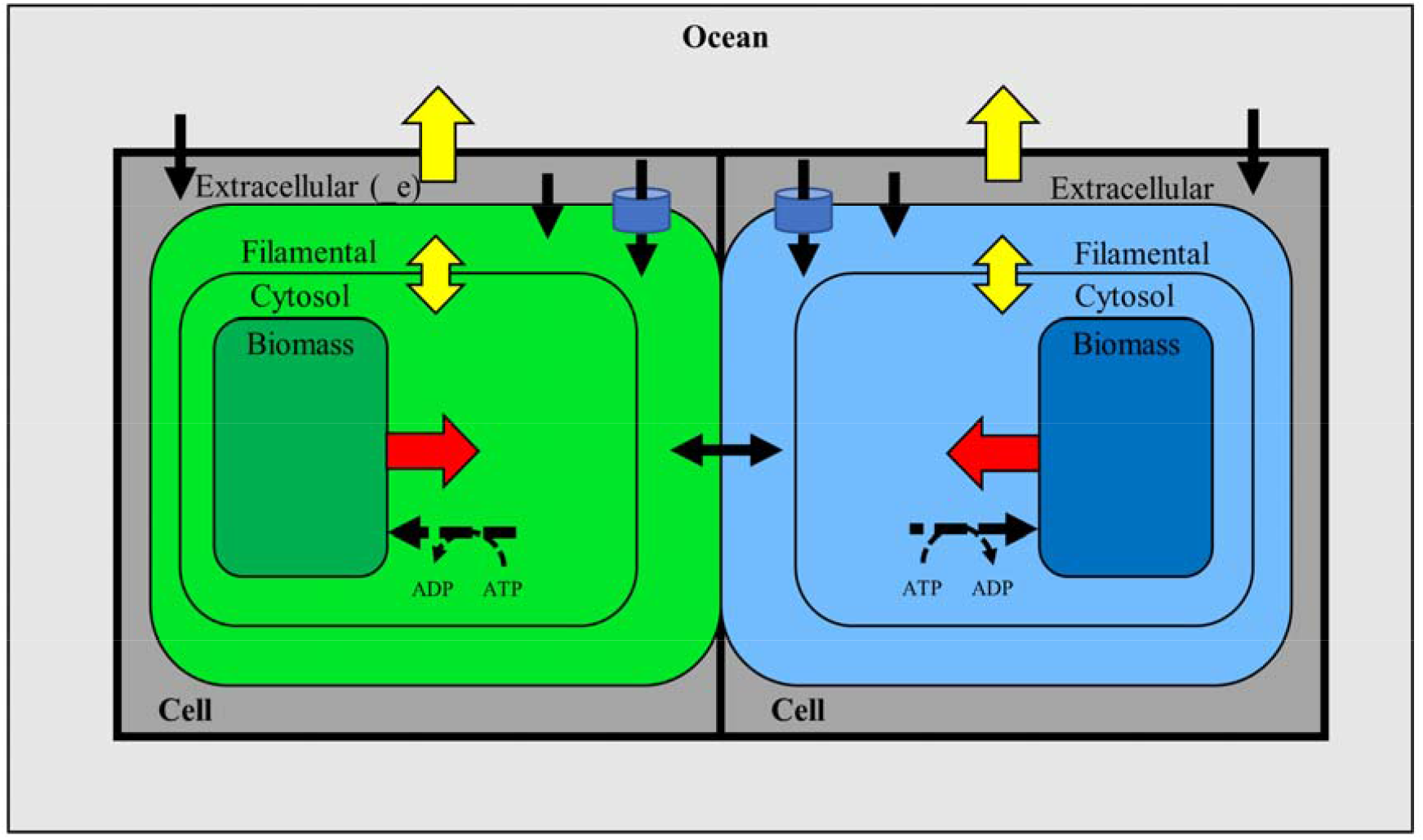
Scheme of transport controls in the agent-based modeling framework between and within cell and ocean agents. Agents (cells and oceans) are indicated in bold lettering and are separated by bold borders. Compartments within are arranged into static biomass (_b), cytosol (_c), filamental (_f), and extracellular (_e). Compartments govern exposure between and within agents. Biomass and cytosol are exposed only to their cell owners. Cytosolic compounds can be converted into filamental compounds through transport mechanisms. Filamental compounds are exposed to other cell members of the filament agent and can be converted into extracellular compounds through transport reactions. Extracellular compounds can be released into ocean (environment) agents or can be taken up from ocean agents. Red arrows imply catabolic transport as a last result; black dashed arrows with ATP/ADP imply metabolic reactions; yellow arrows indicate diffusional transport (such as carbon dioxide or ammonia release); and black arrows imply free or membrane diffusion.

A more comprehensive analysis of the objective equation can be seen in our previous paper (2). Briefly, we considered the primary objective for a cell to create biomass and its primary metabolite in tandem, consistent with a Pareto weighting algorithm described in previous work. The primary objective of biomass (*x*) and metabolite (*m*) can be thus formulated as:

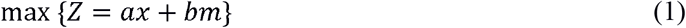

Where *a* and *b* are weights that add up to 1. If this resulted in an infeasible solution, the model progressed with objective function:

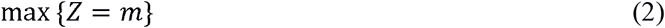

And finally, if that failed, with a catabolic framework where *x*_*c*_ represents the biomass catabolized:

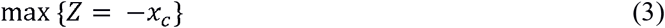

### Rules of Behavior

We improved the MiMoSA modeling framework by the addition of new rules of behavior specific to

*Trichodesmium* and how it handles both oxygen and nitrogen metabolism. Details are provided below.

#### Nitrogenase Inhibition

Three models describing the inhibition/poisoning of nitrogenase by oxygen were evaluated: (i) an exponential model whose parameters were adjusted to fit experimental data, (ii) a logistic model based on the internal oxygen to nitrogen ratio in *Trichodesmium* (28) and (iii) a Michaelis-Menten competitive inhibition model developed for *Azotobacter vinelandii* (29). Details of all three models are below and parameters are provided in Table 1.

**Table 1.** Best fit parameters for modeling nitrogenase inhibition by oxygen.

| Parameter | Value | Units | Model | Source |
| --- | --- | --- | --- | --- |
| k | 3.312 | N/A | Logistic | (30) |
| c | 1.264 | N/A | Logistic | (30) |
| $K_m$ | 0.0488 | mM | Michaelis-Menten | (29) |
| $K_I$ | 0.0182 | mM | Michaelis-Menten | (29) |
| $\beta$ | -1.495 | N/A | Exponential | (30) |
| $\gamma$ | 0.818 | N/A | Exponential | (30) |

Exponential model:

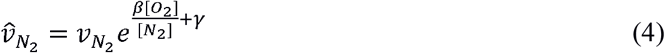

Logistic model:

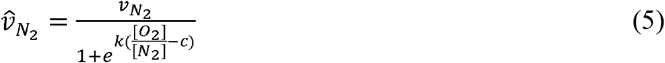

Michaelis-Menten competitive inhibition:

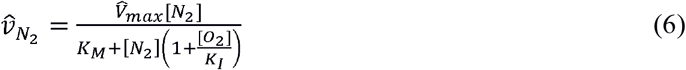

Where 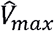 is the flux version of *Vmax* and is calculated as (since *Vmax=f* ([*E*_0_],*k*_*cat*_):

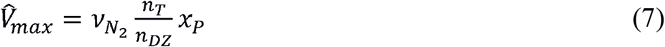

Where 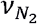 is the unrestricted flux of N_2_ assuming no inhibition or nitrogenase kinetics, *n*_*T*_ is the number of total cells in a filament, *n*_*DZ*_ is the number of diazotrophic cells in the filament, and *x*_*p*_ is a protein- adjustment constant (intended to reflect [E_0_]). Note that 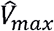 as written is given as a molar rate (mmol (g DW h)^−1^). The ratio of total cells to diazotrophic cells multiplied by the unrestricted flux is meant to resemble the parameter *k*_*cat*_ since nitrogenase only functions in diazotrophic cells. We calculated the parameter *x*_*p*_ as a linear function of biomass accumulation where initial nitrogenase manufacture can be assumed to be a piece of initial biomass development, so it scales linearly with biomass and some experimentally derived parameters:

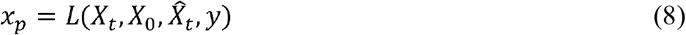

Where *X*_*t*_ is the biomass at time *t, X*_0_is the starting biomass, 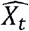 is the biomass at which nitrogenase flux at a maximum, and *y* is the initial nitrogenase present at the beginning of the light period. The final equation is:

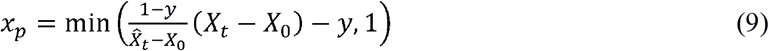

The model was trained on experimental evidence indicating that maximal nitrogenase flux does not occur until 4 hours into the light period (10) likely due to nitrogenase’s inactivity at night as well as the high oxygenation of photoautotrophic low-respiration cells due to carbon deplete conditions. However, it was not trained on the actual value of nitrogenase, simply that there is an initial delay. Since diurnal cycling is out of the scope of this current model iteration, these parameters were lumped into an arbitrary delay term, tying nitrogenase flux to biomass production and allowing the fraction of *V*_*max*_ to reach 100% midway through the cycle under ideal growth conditions assuming a linear relationship between biomass and nitrogenase flux.

#### Oxygen Sequestration

To evaluate the effect of changes in internal oxygen concentrations within the cell and trichome, we evaluated four conditions:

- Anaerobic, which forces all intracellular oxygen to be consumed completely during each time step via respiration reactions
- Fermentative, which allows extra carbon available after all the oxygen is consumed to be directed through common fermentation reactions and the excretion of associated products
- Microaerobic, which restricts intracellular oxygen concentrations to a maximum, described in detail below
- Aerobic, no rules about oxygen concentrations inside the cell were imposed

For each of these conditions, the two major metabolic oxygen sequestration mechanisms were allowed: respiration and Mehler reactions.

For microaerobia, we defined allowable oxygen as the concentration where the nitrogenase activity does not fall below a certain percent activity, α, definited as the ratio of inhibited (*V*) over uninhibited nitrogenase rate (*V’*) at a nitrogen concentration, [*N*_2_] with kinetic parameters (defined above and in the Supplementary Information) 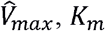, and *K*_*i*_:

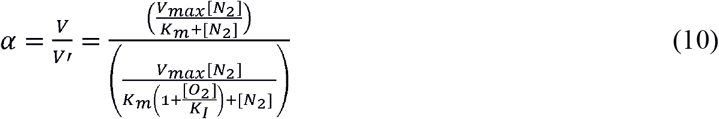

αis defined as 0.9 for this study, so the point at which 90% nitrogen flux is obtainable in the presence of atmospheric O_2_, selected since that is the approximate activity ratio at normal atmospheric conditions according to the literature (30). The “maximum” oxygen concentration [*O*_2_] _*i*_ is:

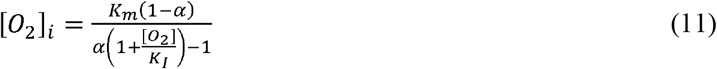

Which implies a Mehler-required flux, ν_*M*_, (divided equally between both Mehler reactions) of:

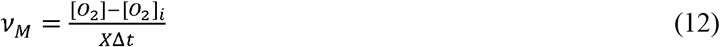

Mehler reactions are only applied to diazotrophs as photoautotrophs do not fix nitrogen. Mehler reactions are downregulated as nitrogenase is downregulated. Therefore, if there is enough reactive nitrogen (ammonium, cyanophycin, urea) to replace N_2_ fixation, Mehler fluxes are switched off according to the algorithms described in Supplementary Files 6-9. Mehler reactions are tethered to nitrogenase flux; the algorithm conditionally chooses whether to require oxygen consumption based on whether nitrogenase flux is required. Nitrogenase flux is required only in the absence of alternative nitrogen sources, which forms a nested condition. Nitrogen acquisition is a condition of cell growth, and therefore this governs – with carbon availability – whether the cell conducts anabolism, catabolism, or stationary production.

#### Photosynthetic Oxygen Evolution Limits

In cases where photoautotrophic oxygen evolution becomes greater than diazotrophic capabilities to feasibly maintain microaerobia via glycolysis or Mehler reactions due to a lack of available substrates, a limit is imposed on photosynthetic oxygen evolution. If nitrogen fixation by diazotrophic cells becomes growth limiting (by dropping below 90% of oxygen free flux), each photoautotrophic cell is limited to producing O_2_ at a rate that is proportional to the previous step’s overall O_2_ consumption corrected for cell ratios 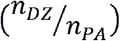 in the filament:

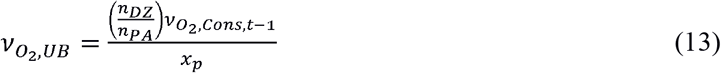

The rationalization for this upper limit is that there is no phenotypic benefit to the cells above where diazotrophs can feasibly consume O_2_, and this rids the model of a futile energy sink as a negative feedback mechanism. Future investigations warrant an investigation into the photosynthesis’ regulation in the presence of high O_2_.

#### Diffusion and Transport of External Nitrogen

We calculated the diffusion of nitrogen sources to cell-specific extracellular compartments (with moles available denoted by *N[e]*) by distributing all moles of a compound in the ocean (environment) gridspace to the extracellular compartment (*N[o]*) divided by the number of other cells in that ocean gridspace (*n*_*c*_). We used an uptake radius factor, *f*_*r*_, cubed to make it per volume:

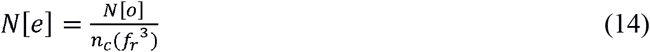

The uptake radius factor, *f*_*r*_, was estimated by fitting the uptake radius over a 12 hour daytime period to match growth characteristics and nitrogen inhibition data at 20 μM NO_3_; this was then evaluated to determine if this value was a good fit for other conditions. As the uptake radius factor grows, the available nitrate for a timestep decreases and leads to less inhibition of nitrogenase. This has been reported in previous literature (28). For gridspaces of 100 μ*m* × 100μ*m* (containing a maximum of 100 cells), we found *f*_*r*_ to be 2.5, allowing an effective radius of 40 μ*m*, constraining to whole-cell widths (see Figure S1). We calculated total nitrogenase flux, *η*as:

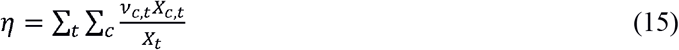

We weighted the individual cell flux for cell *c* at time *t* (ν _*c,t*_) by its individual biomass (*X* _*c,t*_) and dividing by the total cellular biomass at time *t* (*X*_*t*_) and summing over all time steps. We calculated the inhibition percent, ω, under condition indexed *q* similarly to the experimental results (28) as:

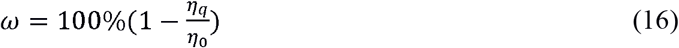

Where η_0_ is the total nitrogenase flux in unamended media.

#### Inhibition of Futile Cycling

Cyanophycin contains both nitrogen and carbon and therefore can be used in the model as a carbon source, but this leads to futile cycling that is likely not physiological. To prevent futile cycling of carbon from cyanophycin in lieu of *de novo* carbon fixation, we implemented a limit on cyanophycin degradation. We did this by limiting the maximal flux of urea and ammonium release using a logistic model because it promoted simple switch-like behavior and has been used for other biological (31) and chemical (32) studies.

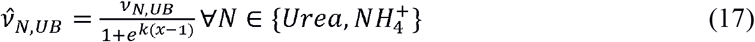

Where 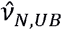 is the allowable maximum output flux, ν _*N,UB*_ is the maximum unrestricted output flux, *k* is a constant estimated as 2ln (9) to allow 90% flux at *x*= 0 and 10% at *x*= 1, where *x* is the measure of a cell’s ability to cover *de novo* nitrogen demands with internal contents, or:

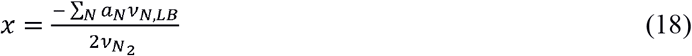

*N* is the set of all reduced nitrogen inside the filament and *a*_*N*_ is the number of nitrogen atoms in one molecule of that substance. It is negative since uptake is negative (*x* will be positive).

In conditions where externally available reduced-nitrogen compounds, *x*_2_, are immediately available, autoinhibition is shut down and 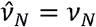 since cells are unlimited by nitrogen in these cases.

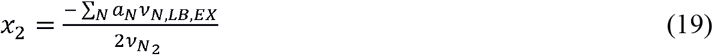

We compared four different models of nitrogen autoinhibition: (i) full autoinhibition by both “free” reactive nitrogen compounds (urea and ammonium), (ii) no autoinhibition of urea or ammonium production, (iii) autoinhibition of only ammonium production, and (iv) autoinhibition of only urea production. Based on comparison with experimental data, we chose to incorporate the full autoinhibition model as it matched most closely cyanophycin accumulation (2) and nitrogen loss to environment (33).

## RESULTS AND DISCUSSION

### Improved Genome-Scale Metabolic Model

The focus of this study was the interplay of nitrogen and oxygen metabolism, so we focused on manually curating the previously published metabolic model (25) to enumerate all the metabolic pathways which metabolize the major building block of nitrogen storage compounds, β-aspartyl arginine (14, 34, 35). The majority of metabolic reactions added to the updated model are for the conversion of arginine into central metabolic intermediates (see Figure 3). *T. erythraeum* has an exceptionally extensive catalog of L-arginine catabolic pathways, perhaps due to the need to recycle stored nitrogen despite the cost of maintaining these pathways (1). To aide in the comparison of these pathways, we have calculated the changes in Gibbs free energy of each pathway shown in Figure 3 and detailed in Table 2 as well as the number of carbon and nitrogen atoms liberated. It is worth noting that these are static estimates of Gibbs free energy, a reasonable assumption given that all of these pathways are close together and the variability in concentration, temperature, pH, or other factors among them is unlikely to be large. From this analysis, it becomes clear that the cell maintains the multitude of pathways to enable the efficient use of stored nitrogen (and by default, carbon) regardless of the energy state of the cell. Cells may also choose pathways based on which central metabolic pathways need supplemental carbon as each pathway results in different metabolites. The updated genome-scale model of *Trichodesmium*, with these updated arginine catabolic reactions is provided as Supplemental Files 2 and 3.

**Figure 3.**
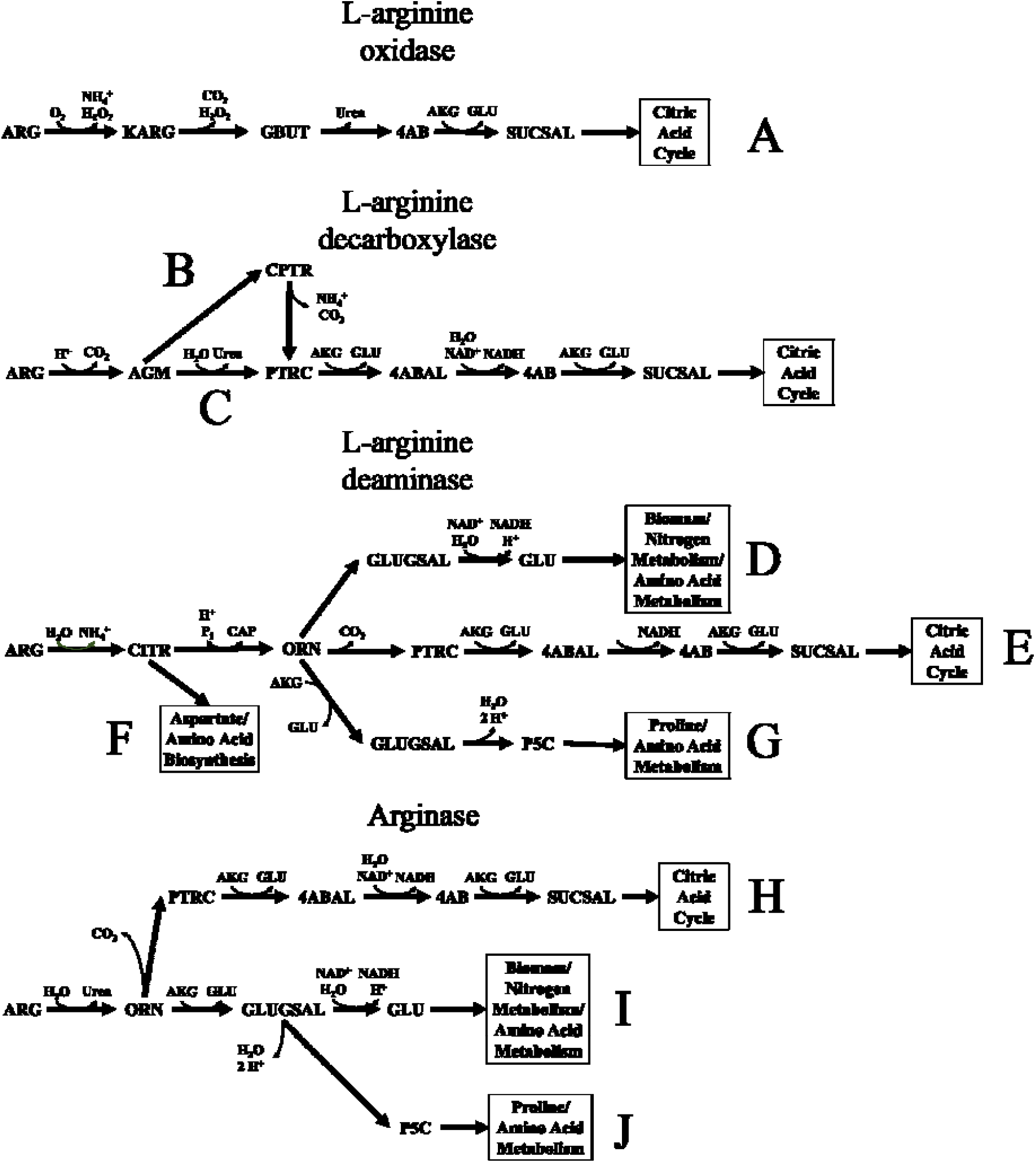
L-arginine catabolism pathways. Each pathway (indexed by letter to Table 2) is a separate metabolic pathway to degrade L-arginine into central metabolic intermediates. Adapted from information in (1). Abbreviations: 4AB: 4-aminobutanoate, 4ABAL: 4-aminobutanal, AGM: agmatine, AKG: -ketoglutarate, ARG: L-arginine, CITR: citrulline, CPTR: N-carbamoylputrescine, GBUT: guanidinobutyrate, GLU: L- glutamate, GLUGSAL: L-glutamate 5-semialdehyde, KARG: ketoarginine, ORN: L-ornithine, P5C: 1- pyrroline-5-carboxylate, PTRC: putrescine, SUCSAL: succinic semialdehyde.

**Table 2.** Energy analysis of arginine catabolism pathways. The sum of Gibbs is the sum of all participating reactions’ Gibbs free energies (36). Gibbs free energy is calculated via the group contribution method (https://pubmed.ncbi.nlm.nih.gov/18645197/) at pH 7.3, ionic strength 0.25, and using the biological standard Gibbs free energy.

| Pathway | Liberated N<br>(mol N) | Liberated C<br>(mol C) | $\sum \Delta G$<br>(kJ/mol) |
| --- | --- | --- | --- |
| A | -3 | -2 | -30.3 |
| B | -2 | -2 | -23.8 |
| C | -2 | -2 | -17.9 |
| D | -1 | 0 | -9.38 |
| E | -1 | -1 | -11.5 |
| F | -1 | 0 | -12.3 |
| G | -1 | 0 | -11.0 |
| H | -2 | -2 | -17.9 |
| I | -2 | -1 | -15.7 |
| J | -2 | -1 | -17.3 |

### Modeling Nitrogenase Inhibition Oxygen Sequestration Methods

It is well documented that nitrogenase activity is negatively impacted by oxygen (37-39); this is especially problematic in nitrogen fixing photosynthetic cells due to the production of oxygen by water splitting. Unlike other diazotrophic cyanobacteria, which separate nitrogenase from oxygen temporally or spatially, *Trichodesmium* trichomes fix carbon and nitrogen simultaneously. To gain insight into how *Trichodemsium* can achieve this, we used the MiMoSA model, which allows us to track nutrients and cells in space and time, to evaluate different models of nitrogenase inhibition as well as oxygen sequestration methods. First, we used three different models (exponential, logistic and competitive inhibition) to model the inhibition/poisoning of nitrogenase by oxygen and compared simulations results to experimental data (10). We found that Michaelis-Menten competitive inhibition resulted in the lowest root mean square error: 0.06 versus 0.103 for logistical model and 0.101 for the exponential. The next step was to determine how the trichome is able to simultaneously carry out nitrogen and carbon fixation without the use of structurally distinct cells (heterocysts).

We tested four cases to determine cellular preferences for oxygen sequestration: (i) anaerobic, which forces all intracellular oxygen to be consumed completely during each time step via respiration reactions, (ii) fermentative, which allows extra carbon available after all the oxygen is consumed to be directed through common fermentation reactions and the excretion of associated products, (iii) microaerobia, which restricts intracellular oxygen concentrations to a maximum determined by the Michaelis-Menten competitive inhibition model (0.368 μM at atmospheric conditions) and (iv) aerobic, where no rules about oxygen concentrations inside the cell were imposed. Selected results of simulations are shown in Figure 4 alongside experimental data collected in our lab for cyanophycin content (2) and published data for nitrogen fixation rates (3). It is important to note here that the cyanophycin data and the initial delay of nitrogenase expression was used originally to train the MiMoSA model (2) but the value of nitrogen fixation rate reported by Kranz et al. was never used for any training and thus serves as a validation data set. These simulations imply that the cell operates in a microaerobic state to balance the need for carbon and nitrogen fixation simultaneously. The accumulation of cyanophycin, the major nitrogen storage compound, actually appears to require oxygen in the cell, as both the fermentative and anaerobic simulations are not capable of accumulating the proper amount of cyanophycin to support nighttime activities in the cell. Since there is a dramatic difference in cellular energy production when respiration is used versus anaerobia, this also seems to be energy driven: the storage of cyanophycin requires a minimum cellular energy level. The different oxygen sequestration methods also appear to play a role in when the peak nitrogenase flux occurs. It appears that the timing of the peak of nitrogenase activity is also a function of oxygen concentration within the cell; fermentative and anaerobic simulations have peaks prior to 4 hours, microaerobic at 4 hours, while the aerobic peak is closer to 6. Several studies have measured peak nitrogenase activity 4 – 6 hours after the onset of light due to the peaks of oxygen within the cell occurring at the onset of light and dark (10, 30, 40). Not only is the microaerobic simulation predicting a peak at 4 hours, it also quantitatively matches the data from Kranz et al. (3), illustrating the power of this modeling approach. When we can track cells and nutrients in space and time, the predictive power of the metabolic model increases dramatically.

**Figure 4.**
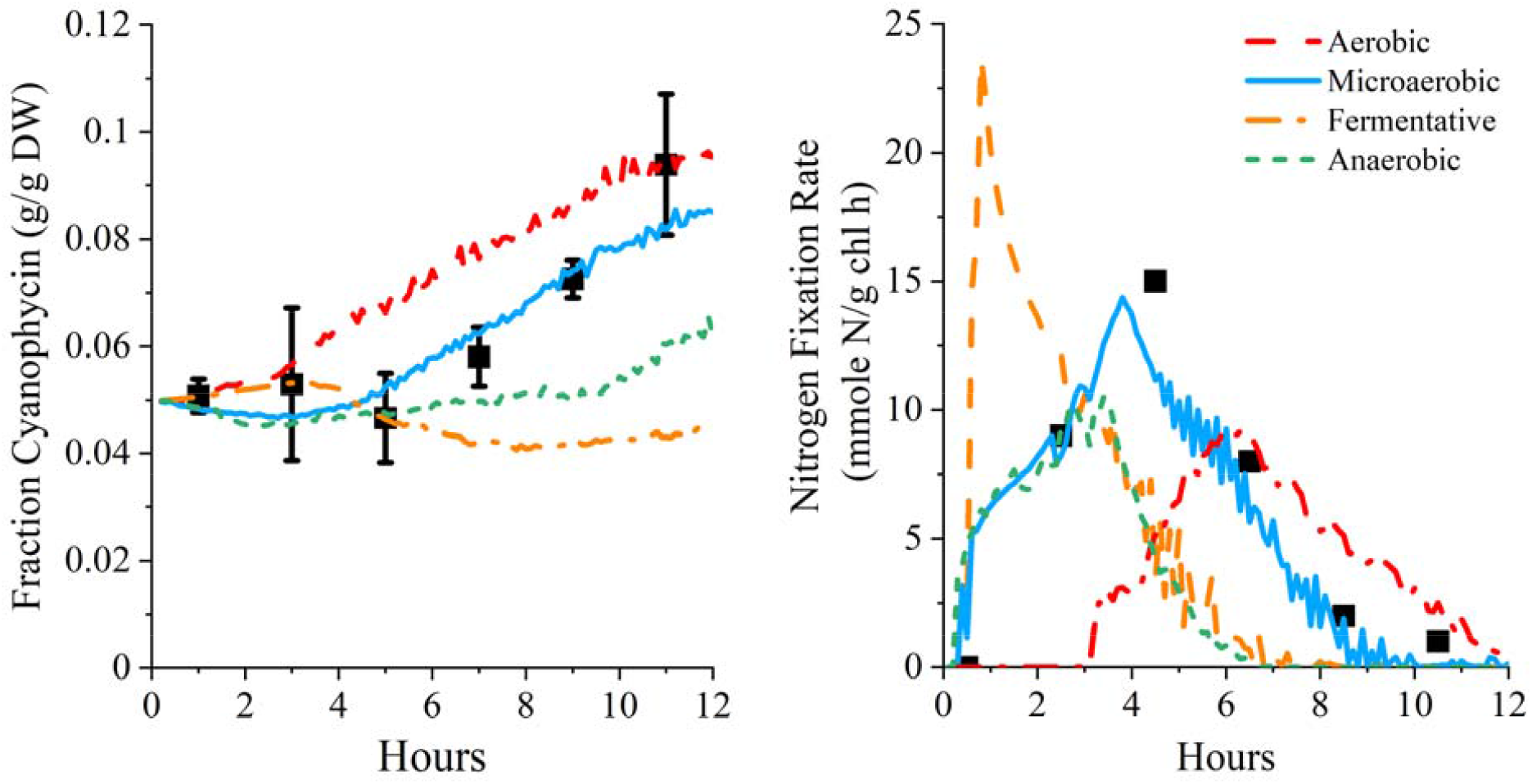
Evaluating potential oxygen sequestration approaches using MiMoSA model. Black squares are experimental data from (2) for cyanophycin content and (3) for nitrogenase fixation rate (interpreted from Figure 2A). From both visual inspection of the data and statistical analysis, microaerobia is the best approach for protecting nitrogenase from oxygen while still allowing respiration to occur for energy production.

### Spatial Effects on Metabolism

Unlike most constraint-based metabolic models, the MiMoSA modeling approach allows us to track individual cells and nutrients in space and time; this means instead of modeling average cells in a well-mixed population, we can investigate how individual cells responds to their changing environment. Trichomes of *Trichodesmium* can vary in length from 3 – 150 cells (7, 41); the geometric mean trichome length reported from *in situ* sampling is 13.2 ± 2.3 cells (41). It has also been reported that each trichome contains 1-3 regularly spaced, non-terminal diazocytes (diazotrophic cell regions) within the filament; the localization of diazocytes in the center of the filament is thought to help protect nitrogenase from oxygen despite the lack of structural elements such as polar plugs or thick cells walls used in heterocysts. We used the model to map the fluxes of crucial carbon, nitrogen and energy metabolic pathways to further investigate the role location within the filament has on metabolism (Figure 5). There are clear differences in how carbon is directed through metabolism in photoautotrophs and diazotrophs; some of these differences arise from differences in constraints, such as the restriction of no photosystem II flux in diazocytes, and others arise from the model and are less intuitive. As expected, carbon fixation fluxes (mainly through RuBisCO) are lower in diazocytes than photoautotrophic cells (Figure 5A) and diazotrophs do maintain lower RuBisCO activity, consistent with experimental observation (42). Interestingly, despite the low intracellular oxygen concentration in diazocytes, it appears they have measurable flux through Phospho*enol*pyruvate carboxylase (PEPCX) and oxaloacetate lyase (OAA Ly) (see Figure 5). These enzymes are known to act in concert in C4 plants as a carbon concentrating mechanism (43, 44). PEPCX is also known to serve a role in regulating TCA cycle flux by refilling oxaloacetate (45), which is also a potential role for PEPCX in the diazocytes, as the TCA cycle is both a mechanism to reduce intracellular oxygen and to produce the energy necessary to fix nitrogen. Diazocytes, as expected, have high higher fluxes through nitrogen assimilation pathways (Figure 5C and D). Cyanophycinase (CPHASE) only functions in photoautotrophs while cyanophycin synthesis (CSyn) and nitrogenase (NITR) occur only in diazotrophs. Urease (UREA) is only functional in diazotrophs, implying that UREA is an essential part of the progressive nitrogen-recycling apparatus. Figure 5D indicates that photoautotrophic cells prefer to degrade cyanophycin using L-aspartate lyase (ASPLS) and arginase (ARGAS), both of which provide intermediates for amino acids and the TCA cycle with minimal carbon loss. These preferences demonstrate some of the utilities of a diverse L-arginine pathway as a carbon-maintenance tool but fail to demonstrate a full use for the extensive arginine metabolic pathways encoded by the *T. erythraeum* genome. Figure 5E and F provide some insight into energy metabolism of the cells. Constraints placed on the model based on experimental evidence restrict the photosystem II (PSII) to zero in diazocytes; it appears that diazocytes compensate for the lack of PSII by increasing flux through PSI as reductive potential is a main constraint on nitrogen fixation. It is important to note that this may be an artifact of the assumptions levied on the system, and merits future investigation It is also important to note here that Mehler reactions seem to be limited to diazocytes, reinforcing yet again the importance of keeping oxygen levels at a minimum to protect nitrogenase. It is clear that the location of cells within a filament plays a major role in their metabolic functions and how *Trichodesmium* is able to carry out nitrogen and carbon fixation simultaneously; the spatial tracking in the MiMoSA framework allows us to investigate these differences.

**Figure 5.**
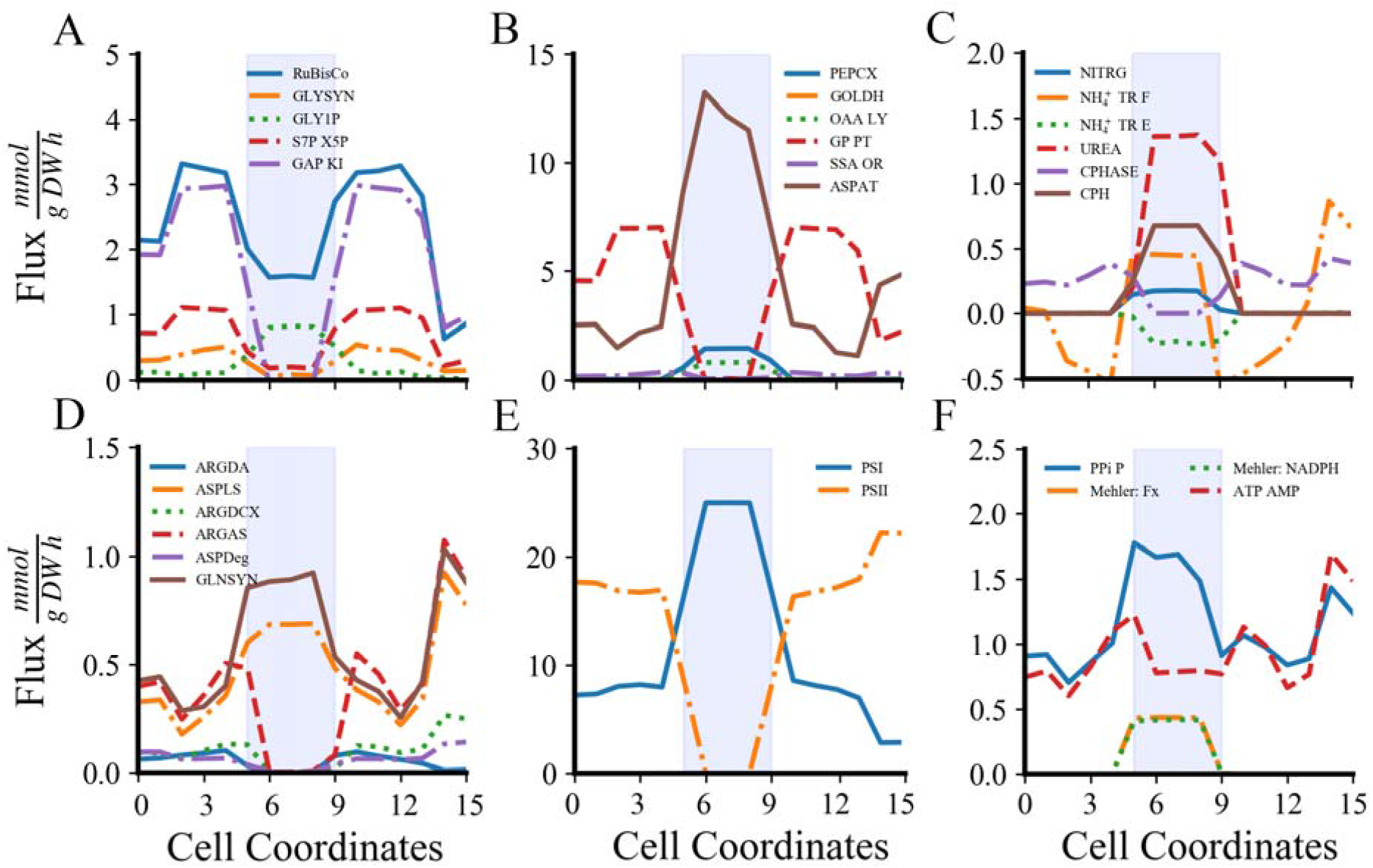
Spatial division of mean metabolic flux during the light period within filaments. Cell coordinates represent the location along filaments in a simulation, where 1 is the cell on one end and 15 is the cell on the opposite end. The shaded area indicates the location of the nitrogen fixing diazocytes. Select reactions of particular interest are highlighted here. Abbreviations: A) RuBisCO: ribulose-1,5-bisphosphate carboxylase; GLYSYN: glycogen synthase; GLY1P: glycogen phosphorylase; S7P X5P: sedoheptulose-7-phosphate: D- glyceraldehyde-3-phosphate transketolase; GAP KI: D-glyceraldehyde-3-phosphate ketol-isomerase. B) PEPCX: phospho*enol*pyruvate carboxylase; GOLDH: glycolate dehydrogenase: NADH; OAA LY: oxaloacetate lyase; GP PT: ATP: 3-phospho-D-glycerate 1-phosphotransferase; Succinate-semialdehyde: NAD^+^ oxidoreductase; ASPAT: L-aspartate 2-oxoglutarate aminotransferase. C) NITR: nitrogenase; NH_4_ ^+^ TR E: ammonium transport from extracellular; NH_4_ ^+^ TR F: filamental ammonium transport, UREA: urease; CPHASE: cyanophycinase; CPH: cyanophycin synthase. D) ARGDA: L-arginine deaminase; ASPLS: L-aspartate lyase; ARGDCX: L-Arginine carboxy-lyase; ARGAS: arginase; ASPDeg: L-aspartate degradation; GLNSYN: glutamine synthase. E) PSI: Photosystem I; PSII: Photosystem II. F) PPi P: pyrophosphate phosphohydrolase; Mehler NADPH: Mehler oxygen consumption with NADPH; Mehler FX: Mehler oxygen consumption with ferredoxin; ATP AMP: ATP:AMP phosphotransferase.

## CONCLUSIONS

Despite the role *Trichodesmium* plays in both the global carbon and nitrogen cycles, not much was known about the exact mechanisms the cell uses to fix both carbon and nitrogen during the day. Here, we used an updated genome-scale metabolic model embedded in the MiMoSA modeling framework (2) to investigate different mechanisms the cell can use to carry out their unique metabolic functions. The main improvement in the updated model is the enumeration of metabolic pathways allowing the degradation of L-arginine (a component of cyanophycin, the main nitrogen storage compound) into central metabolic intermediates and ammonium. This improved model was used to investigate the interplay of nitrogen and oxygen and how *Trichodemsium* is able to operate both metabolic pathways without loss of nitrogenase activity. In order to properly model how nitrogenase is impacted by oxygen, we first used the model to evaluate different models of oxygen inhibition and how closely they matched experimental evidence. Ultimately, in this modeling framework with our ability to track intracellular oxygen concentrations in real time, we determined that a Michealis-Menten Competitive Inhibition model was the best fit. We were also able to evaluate different mechanisms of oxygen sequestration that allows photosynthesis and nitrogen fixation to operate during the day. We compared four different mechanisms: aerobia, microaerobia, fermentative, and anaerobia and found that the mechanism which most closely aligned with the experimental data (2, 3) was microaerobia. The simulations shown in Figure 4, indicate that oxygen is necessary to support the experimentally measured growth rates. By operating in a microaerobic state, the cell is able to balance the need to operate respiratory metabolism, such as the TCA cycle, to generate the energy required to fix nitrogen with the need to protect nitrogenase from oxygen poisoning. Finally, we also took advantage of the spatial resolution and single cell tracking of the MiMoSA modeling framework to examine how location along the filament impacts metabolism. A distinct shift in metabolism is seen moving from the outside of the filament toward the center, potentially due to the difference in nutrient availability. In these simulations, diazocytes operate very similarly, we would expect if the trichome is longer and has more diazocytes, their metabolisms would vary more as diffusion of glycogen in and nitrogen compounds out will start to impact each cell differently. We have shown here that the MiMoSA modeling framework is a powerful tool to simulate growth of more complex mixtures of cells than traditional constraint-based steady state modeling and that the ability to track nutrients and cells in space and time allows us to investigate how these more complex mixtures behave and thrive in different environments.

## Acknowledgements

This work was supported by a grant from the Department of Energy Office of Science, Biological and Environmental Research (BER) Early Career Program grant no. DE-SC0019171 to NRB.

## Contributions

JJG, BMSH and NRB designed the research. JJG performed the research. JJG and NRB analyzed the data. JJG, BMSH and NRB wrote the manuscript. JJG, BMSH and NRB edited and revised the manuscript.

