## Supplementary material for "Investigating the unique ability of *Trichodesmium* to fix carbon and nitrogen simultaneously using MiMoSA": Supplememtary Files

**Description of supplementary files provided with manuscript:**

Supplemental File 1: Updated reactions, genes, and metabolites for this iteration of the genome-scale metabolic reconstruction (.xlsx).

Supplemental File 2: Complete genome-scale metabolic network reconstruction for *Trichodesmium erythraeum* for each cell type, solved in training data for 80 $\mu$E (.json).*

Supplemental File 3: Complete genome-scale metabolic network reconstruction for *Trichodesmium erythraeum* for each cell type, allowing for catabolic digestion of metabolism at 80 $\mu$E (.json).*

Supplemental File 4: Decision algorithm for metabolic regime (.pdf).

**Algorithm 1: Decision algorithm for metabolic regime**

This file describes the decision process for solving cellular metabolic states given their spatiotemporal positions. First, a prospective objective function (*R­_obj_*) is generated for the cell of type τ. A constrained FBA model is created as well as a backflow catabolic model. If a solution exists for the anabolic model (supposing that a cell works first in its own self-interest), the model returns the anabolic solution (see later algorithms). Otherwise, the cell then attempts to support its neighbors by creating compounds specific to its cell type. Finally, it attempts to catabolize sufficient biomass to survive (using the backflow metabolic model). If it fails this, it dies. This methodology follows the logic of other staged multi-objective algorithms like OptCom following the adjustment of scalarized objective parameters along the Pareto Front.

Supplemental File 5: Algorithm for anabolic metabolic optimization (.pdf).

**Algorithm 2: Anabolism**

This file describes the algorithm for anabolism via subsequent relaxation of bounds. First, the simple solution is attempted given available resources from metabolite transport along the filament. If that is infeasible, constraints are relaxed to ignore physiological requirements for waste (O_2_) removal via Mehler’s reactions. If infeasibility persists, the cell is allowed to consume some of its stored (not freely transported) metabolite specific to cell type to achieve anabolism ($\beta$-aspartyl arginine for photoautotrophs and sucrose/glucose dimers for diazotrophs). Given a feasible solution, the other objective reactions represented in the scalarized objective equation are optimized to correct for non-dominant relaxation solutions.

Supplemental File 6: Algorithm for metabolic optimization of primary production (.pdf).

**Algorithm 3: Primary Production**

Given that no biomass can be feasibly made within the relaxation regimes, the cell supposes it can produce its cell-type specific metabolite. If it can, the cell returns that solution. If not, it progresses through relaxation by allowing it to consume its transacted metabolite ($\beta$-aspartyl arginine for photoautotrophs and sucrose/glucose dimers for diazotrophs). If that fails, it maximizes the underlying process (nitrogen fixation for diazotrophs and photosynthesis for photoautotrophs). Failing those, the cell moves to catabolism.

Supplemental File 7: Algorithm for metabolic optimization of catabolism (.pdf).

**Algorithm 4: Catabolism**

The final grasp at life for cells is to recite “Tomorrow and tomorrow and tomorrow” from MacBeth while activating the backflow catabolism model. This is the stoichiometric model that minimizes the number of reactions required to resolve the “biomass” summary metabolite into its components and further into central metabolism. If a solution is found, the consumption of biomass elements are minimized to minimize futile growth.

Supplemental File 8: Abbreviations and definitions of reactions visualized in this study (.xlsx).

Supplemental File 9: Transport reactions added to model for access to local nutrients (.xlsx).

Supplemental File 10: Complete Python optimization framework (.zip with .py).*

Supplemental File 11: Agent-Based Model for use in Repast Simphony in Java (.zip for Repast Simphony).*

* These two files were not compatible with the supplemental file upload so they will be supplied via github when the manuscript is published.

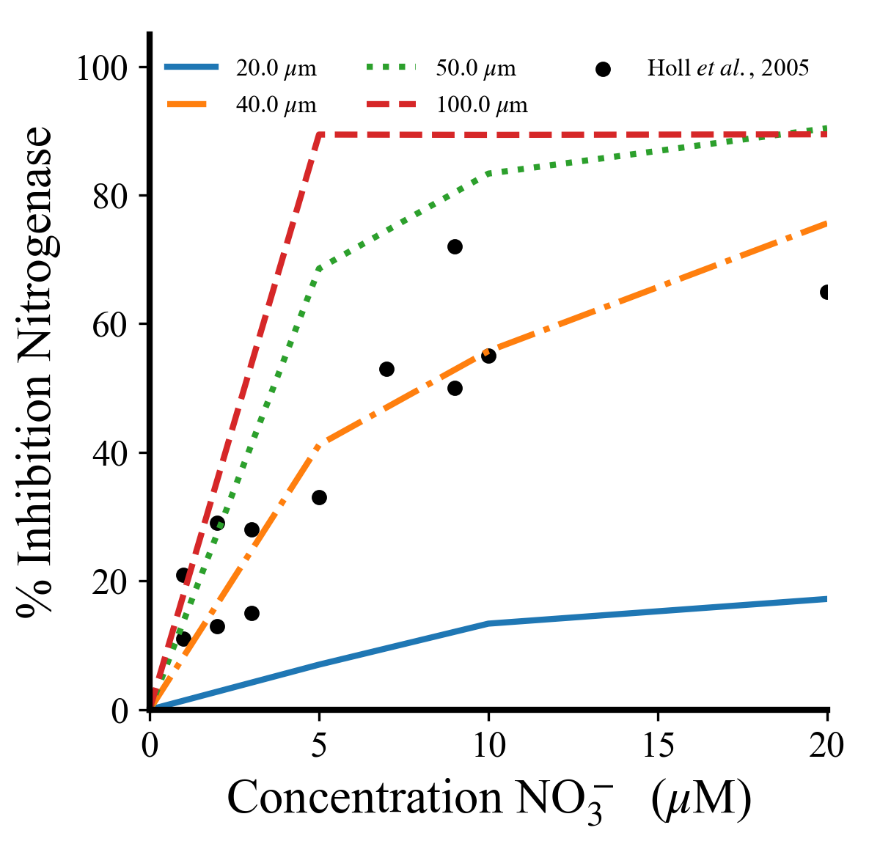

**Figure S1. Comparison of different uptakes radii for nitrate with published data (1).**
