## Supplementary material for "Investigating the unique ability of *Trichodesmium* to fix carbon and nitrogen simultaneously using MiMoSA": Supplemental File 4 Decision algorithm for metabolic regime.pdf

**Algorithm 1:** Metabolic Optimization

**Data:** Genome Scale Model **S**, Flux Constraints $\mathcal{C}$, Objective Weights $\mathcal{W}$, Cell Type $\tau$, Biomass $\mathcal{B}$

**Result:** Optimum **z**, Cellular Fluxes **v**

$R_{obj} = \sum \mathcal{W}^T \cdot \mathbf{v}$

$\mathrm{v}_{exp}$ = Get Export($\tau$)

$\mathrm{v}_{fix}$ = Get Fixation Reaction($\tau$)

$\mathbf{S}_{cat}$ = Generate Catabolic Model(**S**)

**z, v** = Anabolize(**S**, $\mathcal{C}$, $\mathcal{B}$, $\mathcal{B}_T$, $R_{obj}$, $\mathcal{O}$)

**if** *Anabolism successful* **then**

  **return z, v**

**else**

  **z, v** = Primary Produce(**S**, $\mathcal{C}$, $\mathcal{B}$, $\mathcal{B}_T$, $\mathrm{v}_{exp}$, $\mathrm{v}_{fix}$, $R_{obj}$)

  **if** *Primary Production successful* **then**

    **return z, v**

  **else**

    **z, v** = Catabolize($\mathbf{S}_{cat}$, $\mathcal{C}$, $\mathcal{B}$, $R_{obj}$)

    **if** *Catabolism successful* **then**

      **return z, v**

    **else**

      Die

      **return** $\{\}$
