## Supplementary material for "Investigating the unique ability of *Trichodesmium* to fix carbon and nitrogen simultaneously using MiMoSA": Supplemental File 5 Algorithm for anabolic metabolic optimization.pdf

**Algorithm 2:** Anabolism

**Function** `Anabolize` (***S***, $\mathcal{C}$, $\mathcal{B}$, $\mathcal{B}_T$, $R_{obj}$, $\mathcal{O}$)**:**

z = maximize $R_{obj}$
subject to
**S**·**v** = 0
$y_{i,lb} \leq v_i \leq y_{i,ub} \quad \forall i \in \mathcal{C}$
$0 \leq v_j \leq 1000 \quad \forall j \in \mathcal{B}$
**if** $z \geq 0$ **then**
  **z** ← z
**else**
  z = maximize $R_{obj}$
  subject to
  **S**·**v** = 0
  $y_{i,lb} \leq v_i \leq y_{i,ub} \quad \forall i \in \mathcal{C} \setminus \mathcal{M}$
  $0 \leq v_j \leq 1000 \quad \forall j \in \mathcal{B}$
  **if** $z \geq 0$ **then**
    **z** ← z
  **else**
    z = maximize $R_{obj}$
    subject to
    **S**·**v** = 0
    $y_{i,lb} \leq v_i \leq y_{i,ub} \quad \forall i \in \mathcal{C} \setminus \mathcal{M}$
    $0 \leq v_j \leq 1000 \quad \forall j \in \mathcal{B} \setminus \mathcal{B}_T$
    $y_{k,lb} \leq v_k \leq 1000 \quad \forall k \in \mathcal{B}_T$
    **if** $z \geq 0$ **then**
      **z** ← z
    **else**
      **return** failure
**for** $i$ ← 0 **to** *len*($\mathcal{O}$) **do**
  $\hat{z}$ = maximize $\mathcal{O}_i$
  subject to
  **S**·**v** = 0
  $v_{j,lb} \leq v \leq v_{j,ub} \forall j \in$ **v**
  $R_{obj}$ = z
  $v_{obj,k} = \hat{z}_{obj,k} \forall k < i$
**return z**, **v**
