## Supplementary material for "Investigating the unique ability of *Trichodesmium* to fix carbon and nitrogen simultaneously using MiMoSA": Supplemental File 6 Algorithm for metabolic optimization of primary productivity.pdf

**Algorithm 3:** Primary Production

```
Function Primary Production(S, C, B, B_T, v_exp, v_fix, R_obj):
    z = maximize v_exp
    subject to
        S·v = 0    R_obj = 0
        y_{i,lb} ≤ v_i ≤ y_{i,ub}   ∀i ∈ C
        0 ≤ v_j ≤ 1000   ∀j ∈ B
    if z ≥ 0 then
        z ← z
    else
        z = maximize v_exp
        subject to
            S·v = 0
            R_obj = 0
            y_{i,lb} ≤ v_i ≤ y_{i,ub}   ∀i ∈ C
            0 ≤ v_j ≤ 1000   ∀j ∈ B \ B_T
        if z ≥ 0 then
            z ← z
        else
            z = maximize v_fix
            subject to
                S·v = 0
                R_obj = 0
                y_{i,lb} ≤ v_i ≤ y_{i,ub}   ∀i ∈ C
                0 ≤ v_j ≤ 1000   ∀j ∈ B
            if z ≥ 0 then
                z ← z
            else
                return failure
    return 0, v
```
