## Supplementary material for "Investigating the unique ability of *Trichodesmium* to fix carbon and nitrogen simultaneously using MiMoSA": Supplemental File 7 Algorithm for metabolic optimization of catabolism.pdf

**Algorithm 4:** Catabolism

**Function** `Catabolize`(***S***, $\mathcal{C}$, $\mathcal{B}$, $R_{obj}$, $v_{exp}$)**:**

z = maximize $v_{exp}$

subject to

**S**·**v** = 0

$R_{obj} = 0$

$y_{i,lb} \leq v_i \leq y_{i,ub} \quad \forall i \in \mathcal{C}$

$v_{j,lb} \leq v_j \leq 1000 \quad \forall j \in \mathcal{B}$

**if** $z \geq 0$ **then**

**z** ← z

**for** $i \leftarrow 0$ **to** *len*($\mathcal{B}$) **do**

$\hat{z}_i$ = minimize $\mathcal{B}_i$ subject to

$v_{exp} = \mathbf{z}$

$\mathcal{B}_j = \hat{z}_j \ \forall j < i$

**S**·**v** = 0

$R_{obj} = 0$

$y_{i,lb} \leq v_i \leq y_{i,ub} \quad \forall i \in \mathcal{C}$

$v_{j,lb} \leq v_j \leq 1000 \quad \forall j \in \mathcal{B}$

**else**

**return** failure

**return** 0, **v**
