## Supplementary figures and images for "Investigating the unique ability of *Trichodesmium* to fix carbon and nitrogen simultaneously using MiMoSA"

### Figure S1 Scheme of transport controls in the agent-based modeling framework.tif

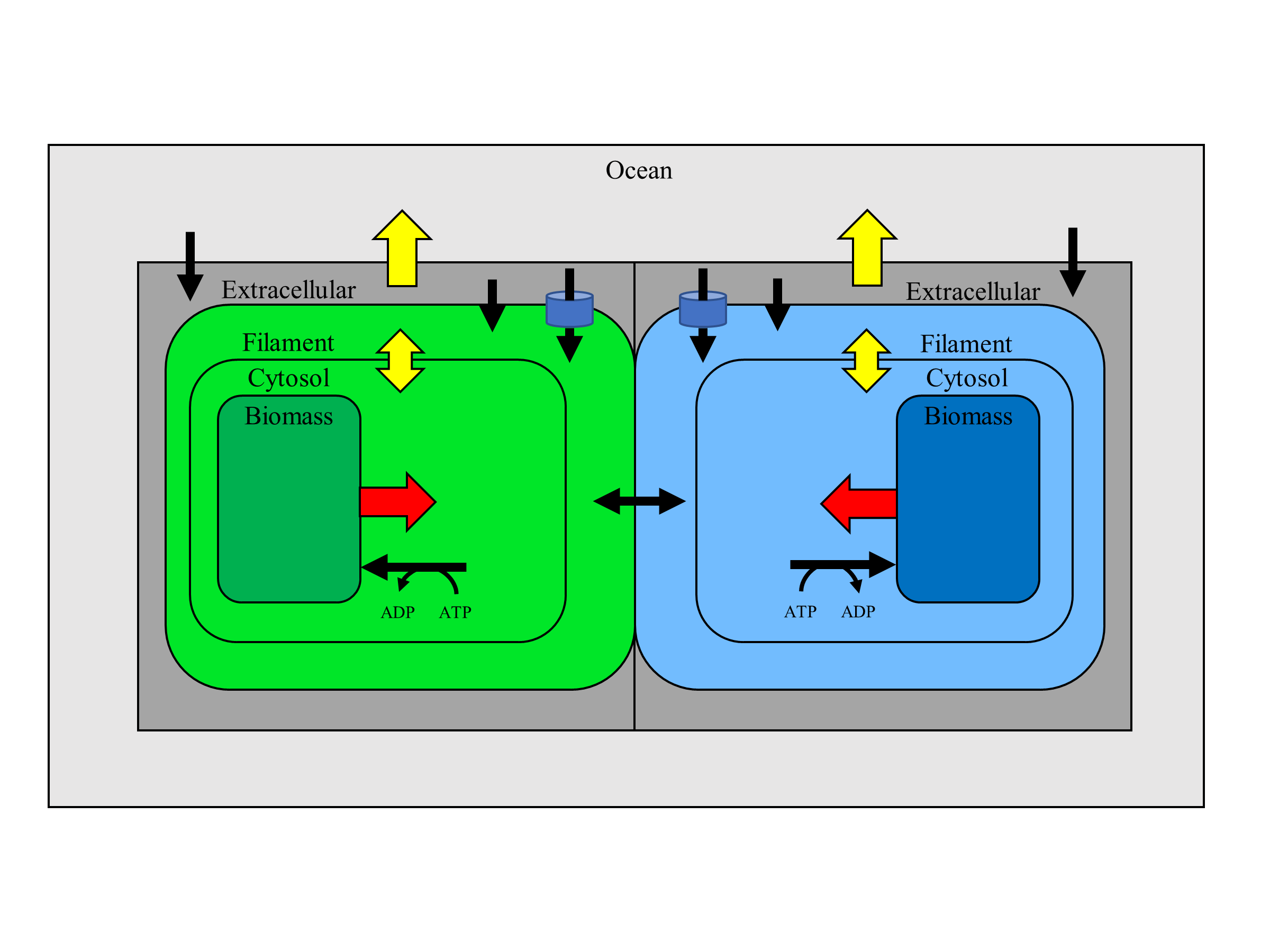

### Figure S2 Contrasting models for nitrogenase inhibition.tif

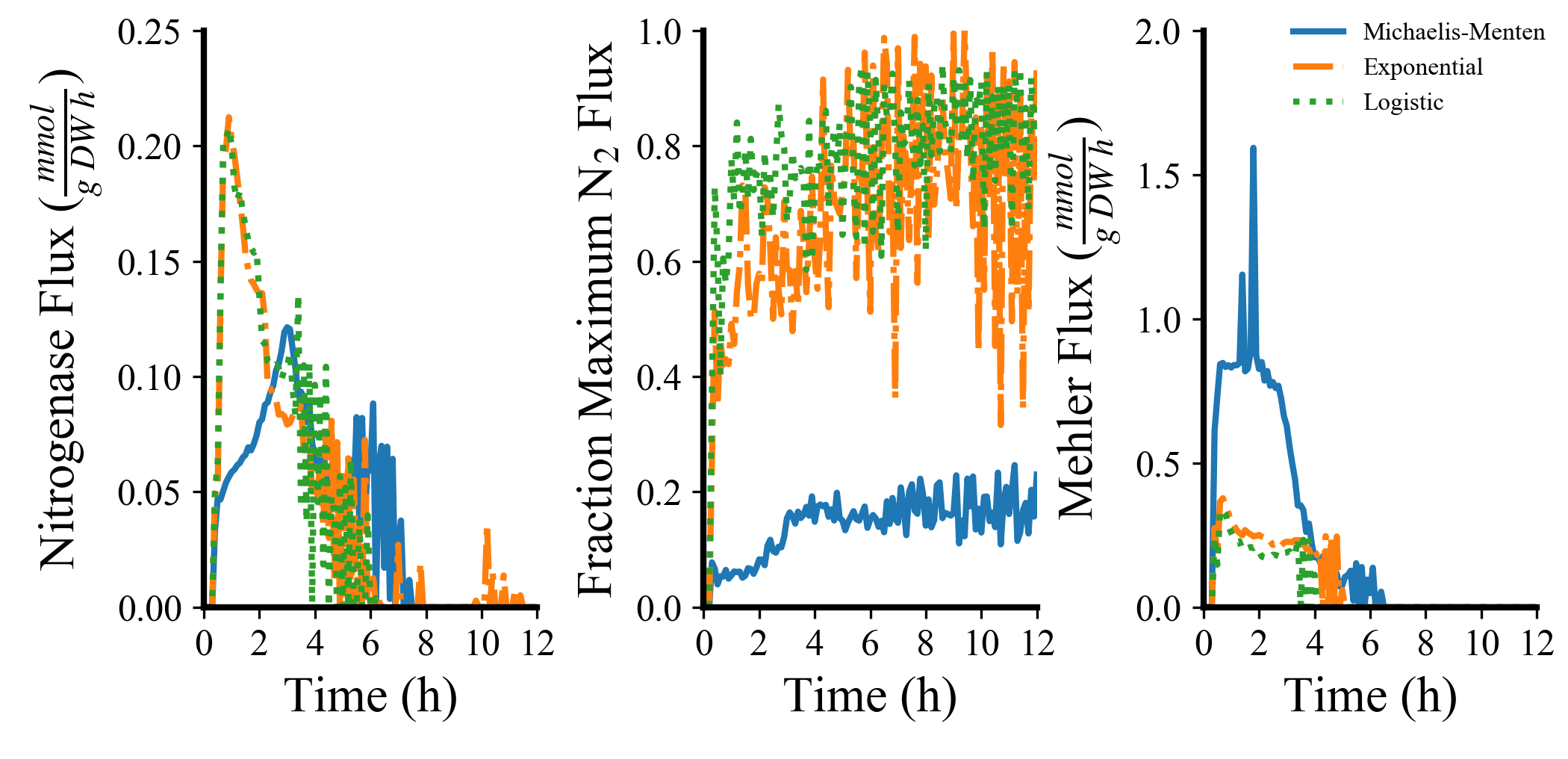

### Figure S4 Prediction of optimal uptake radius through nitrogenase inhibition by nitrate.tif

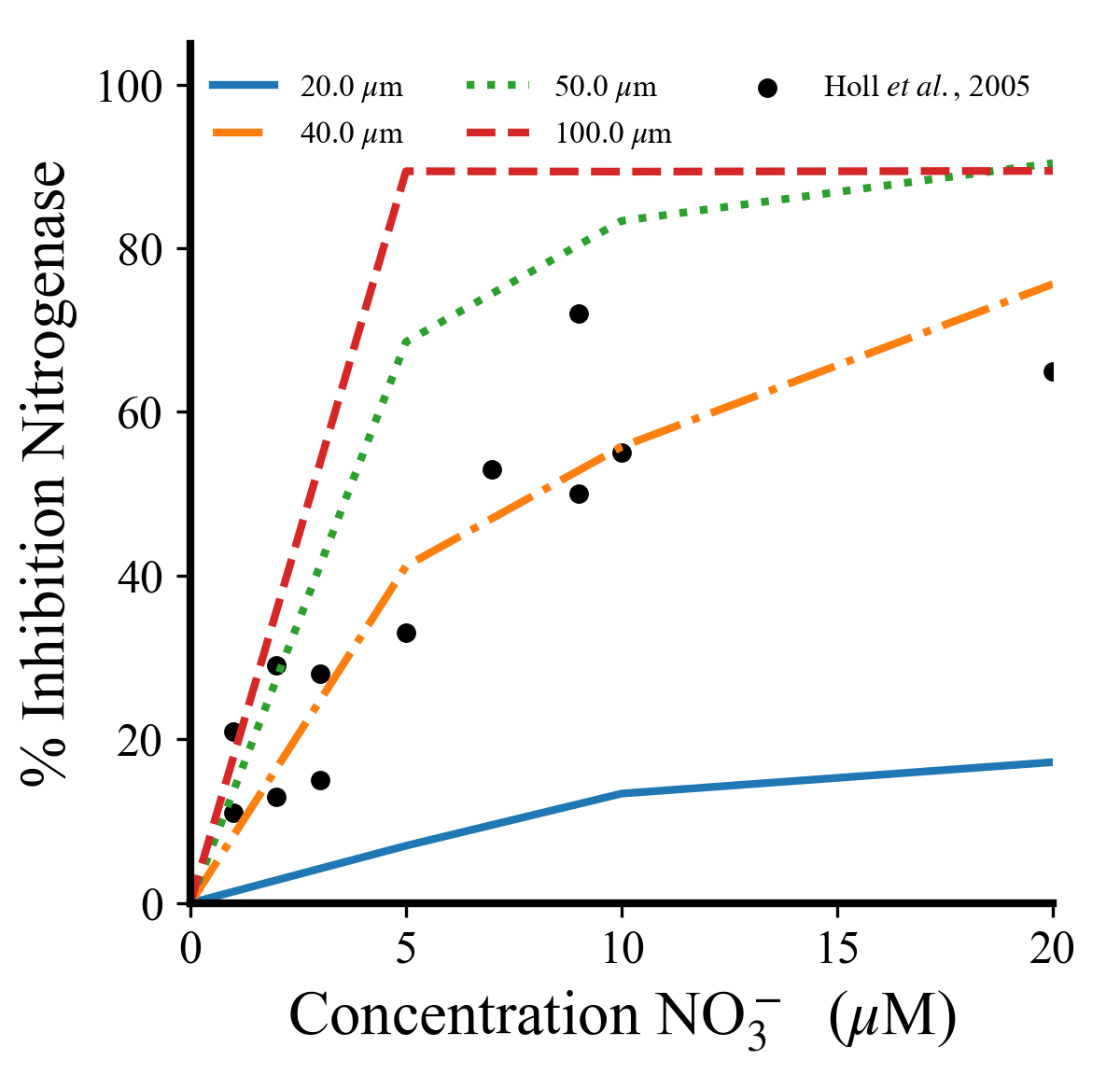

### Figure S5 Effects of nitrogenase delay on nitrogen fixation and biomass outcome.tif

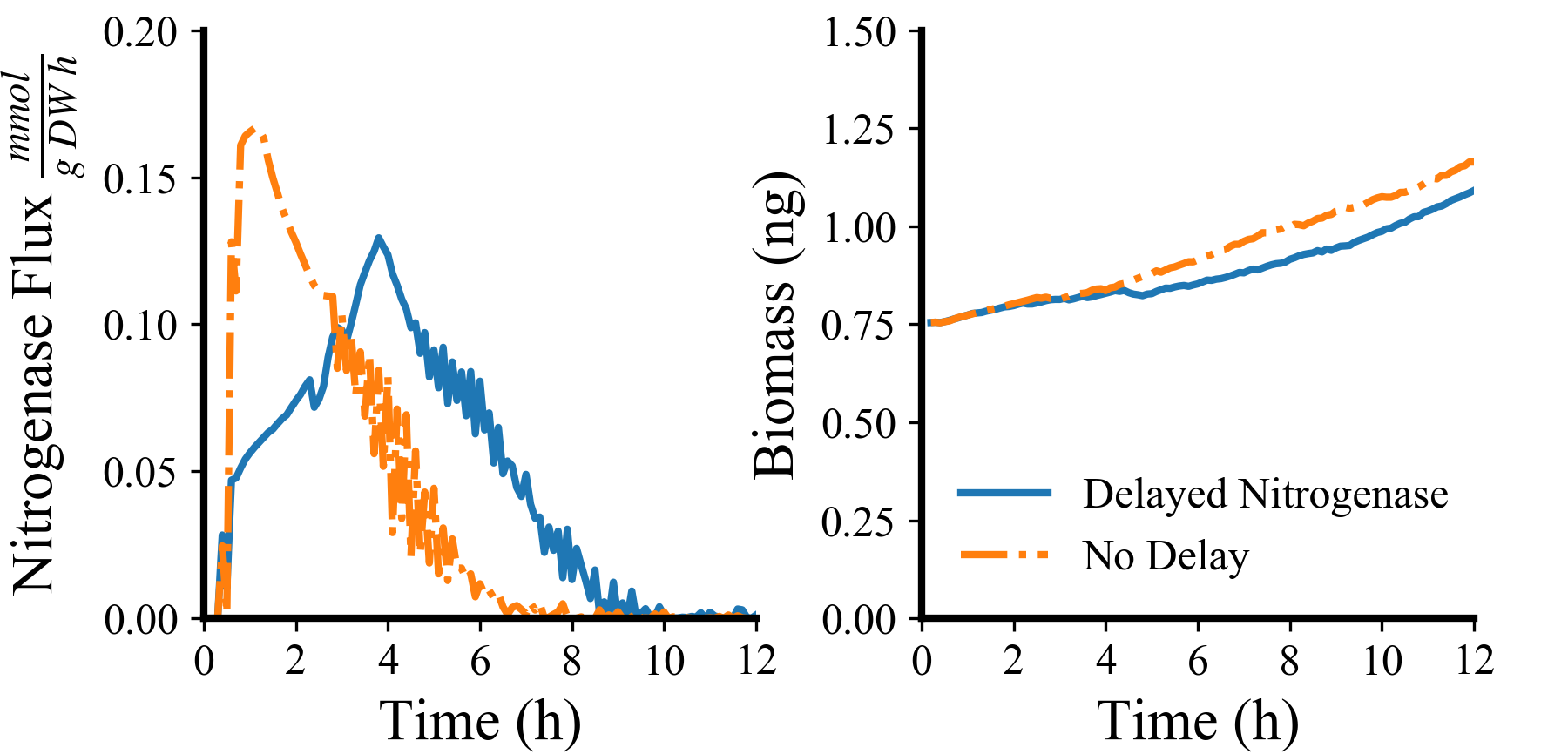

### Figure S6 Temporal flux distributions between cell types in nitrogen metabolism.tif

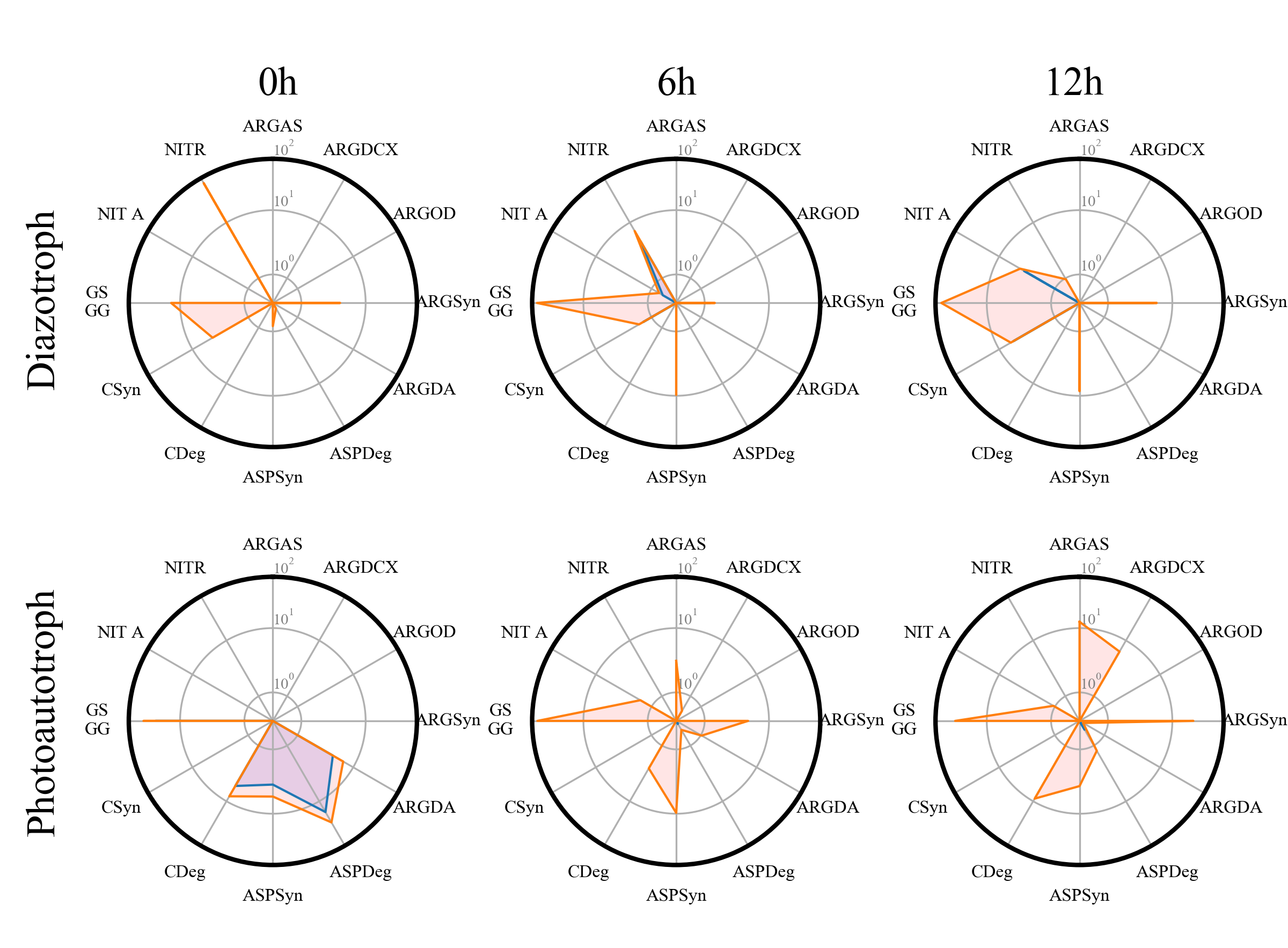

### Figure S7 Nitrogen metabolism distribution in nitrogen amended media.tif

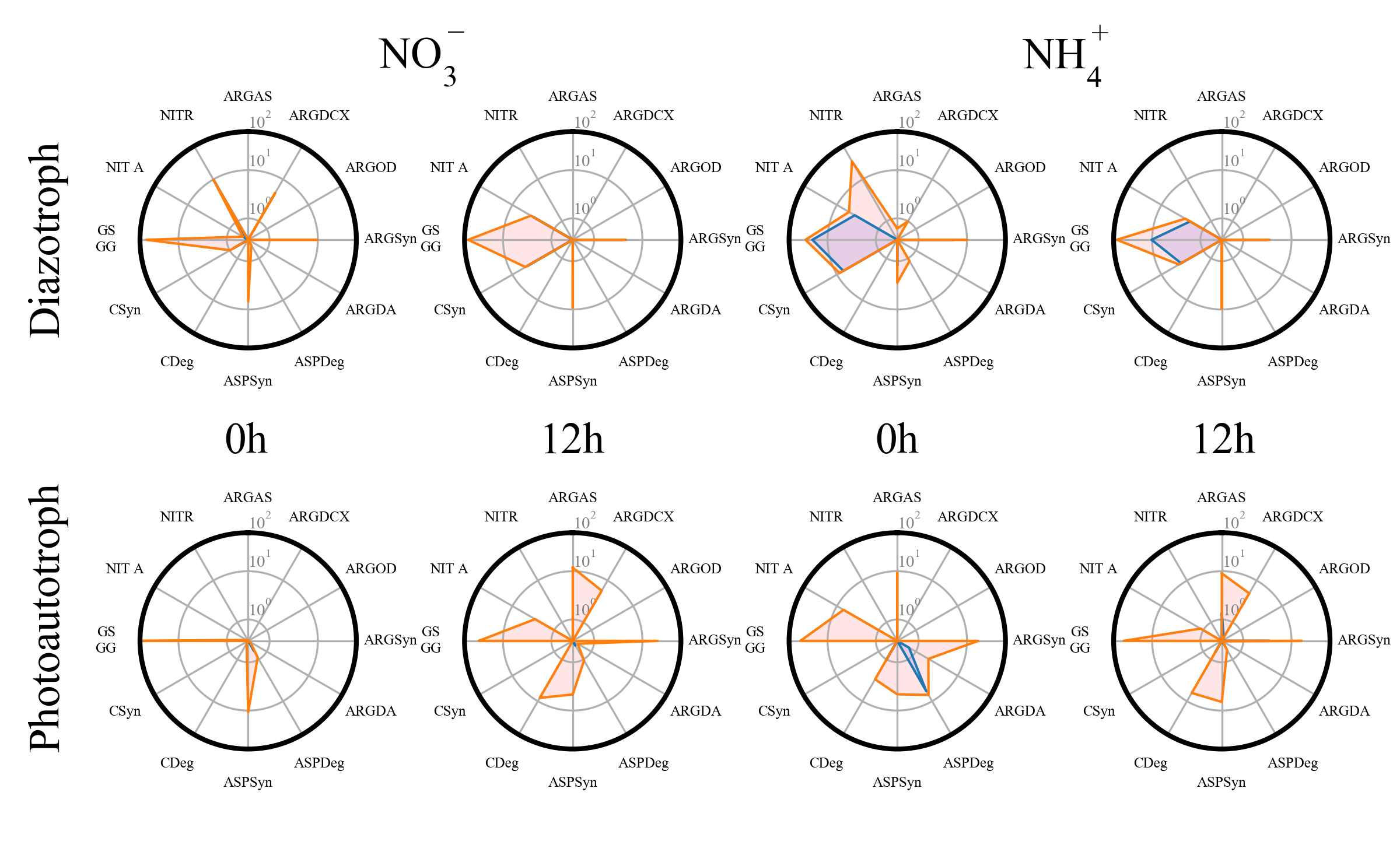
